# Bergmann’s rule in *Microlophus* lizards: testing for latitudinal and climatic gradients of body size

**DOI:** 10.1101/2022.01.18.476846

**Authors:** Ken S. Toyama, Christopher K. Boccia

## Abstract

Broadscale environmental gradients exert selection pressures that can result in macroevolutionary patterns of phenotypic diversity. Bergmann’s rule, one of the most studied ecogeographical patterns, states that among related species or populations, larger body sizes should occur at higher latitudes or colder climates given the efficacy of larger sizes to conserve heat. However, there is still much debate regarding the mechanisms that underlie the rule and whether they should be different between endotherms and ectotherms. Here we test for Bergmann’s rule in *Microlophus* lizards, a genus distributed in western South America and the Galápagos Islands, and test for associations between body size and climatic variables. Our results indicate that mainland *Microlophus* species follow Bergmann’s rule, meaning that larger species are found towards the southernmost latitudes encompassed by the genus’ distribution. This relationship was similar for males and females, suggesting that patterns of sexual size dimorphism are not driven by differing latitudinal gradients of body size between the sexes. We identified negative associations between body size and temperature, precipitation, and isothermality, meaning that larger species are found in colder, drier, and more thermally variable environments. We discuss the possible mechanisms behind this pattern, the implications of insularity and ecology for the evolution of body size in this genus, and emphasize the need for additional natural history information to allow for the elucidation of the mechanisms behind the patterns presented in this work.

## INTRODUCTION

Environmental factors can mediate patterns of phenotypic diversity in nature (Hawkins et al., 2003; Harrison et al., 2020). For instance, geographic variation in climatic variables can produce gradients of selection that ultimately produce systematic phenotypic variation at a macroevolutionary scale (e.g., Chen et al., 2013; Pincheira-Donoso et al., 2019). One of the most well-studied macroecological patterns relating climatic factors and phenotypic traits is Bergmann’s rule, which states that within a lineage, larger species should be found at higher latitudes or colder climates (Bergmann, 1847; Meiri, 2011). Since its formulation, the rule has been confirmed in many taxa and is widely generalizable for endotherms, including mammals and birds (e.g., Ashton et al., 2000; Ashton, 2002; Blackburn and Hawkins, 2004; Meiri and Dayan, 2003). The heat retention hypothesis originally put forward by Bergmann himself (i.e., better heat conservation is achieved at larger sizes given an increase in the volume/surface area ratio) has traditionally been proffered as the main mechanism behind Bergmann’s rule (Meiri, 2011). However, several other mechanisms are thought to play a role in driving the relationships found between body size and latitude across taxa (e.g., Cushman et al., 1993; Atkinson and Sibly, 1997; Ashton et al., 2000; Walters and Hassall, 2006; Wolverton et al., 2009).

For ectotherms, the evidence for Bergmann’s rule is inconclusive. Although several studies have found that ectotherm groups follow the rule (e.g., Ashton and Feldman, 2003; Cruz et al., 2005), several others found no support for any pattern (e.g., Belk and Houston, 2002; Adams and Church, 2008; Pincheira-Donoso et al., 2008) or detected the inverse trend (a decrease in size towards higher latitudes, e.g., Mousseau, 1997; Ashton and Feldman, 2003). An inverse trend is a reasonable alternative expectation for ectotherms given the original heat conservation mechanism, which is based on the maintenance of a preferred body temperature. In contrast to endotherms, ectotherms do not produce heat internally so selection might act on the capacity of the body to gain heat rather than to retain it. Therefore, a small body size might evolve in cooler climates as a consequence of selection for a lower volume/surface area ratio, which enables more rapid heating (Stevenson, 1985). Nonetheless, other mechanisms could be at play for ectotherm groups that follow Bergmann’s rule. For example, hypotheses based on seasonality suggest that larger individuals are better suited to survive longer periods of food shortage given their increased capacity to store fat (Murphy, 1985). Higher latitude environments typically present frequent periods of food scarcity due to seasonality; thus a relationship between body size and seasonality could result in patterns similar to Bergmann’s rule for both endotherms and ectotherms. Similarly, oxygen limitation has been proposed to drive negative size – temperature relationships among ectotherms with aquatic respiration, as oxygen solubility in water decreases at higher temperatures (Rollinson and Rowe, 2018).

Regardless of the mechanism, a relationship between size and latitude driven by climatic factors likely affects other macroevolutionary patterns that are related to size. For instance, Blanckenhorn et al (2006) showed how intraspecific patterns of sexual size dimorphism arise as a consequence of different size – temperature relationships between sexes. Specifically, they showed that latitudinal gradients of male body size were steeper than those of females for a majority of species in their analysis, yielding a relationship between latitude and male-biased sexual size dimorphism that was consistent with Rensch’s rule (i.e. the increase of sexual size dimorphism with size in species where males are the larger sex; Rensch, 1950). Similarly, Tarr et al (2019) proposed that the latitudinal gradients of sexual size dimorphism they observed in Central and North American lizards were predominantly driven by the effect of sexual selection on male body size, which intensified at higher temperatures and lower latitudes. This implies that macroevolutionary patterns like Rensch’s rule might arise as a consequence of underlying processes driving the evolution of body size, such as the mechanisms underpinning Bergmann’s rule.

*Microlophus* is a genus of diurnal and omnivorous iguanian lizards that includes 13 continental species distributed along the western coast of South America (Figure 1) and a radiation of 10 species within the Galápagos archipelago. All species show male-biased sexual size dimorphism, with maximum snout-vent lengths (SVL) ranging from 80 to 155 mm in males and from 65 to 116 mm in females (Toyama and Boccia, 2021). On the mainland, *Microlophus* species span ∼30 degrees of latitude and inhabit different ecosystems including dry forests, intertidal rocky shores and seasonal oases (Dixon and Wright, 1975; Quispitúpac and Pérez, 2008). Most species, with one exception, are found in the coastal regions across the Sechura and Atacama deserts (Dixon and Wright, 1975). *Microlophus arenarius* is the only species found in the inter-Andean valleys of Northern Peru (Dixon and Wright, 1975; Koch et al., 2018; Figure 1B), and likely experiences different climatic conditions compared to other *Microlophus* species (e.g. higher precipitation, Duellman and Pramuk, 1999).

**Figure 1.**
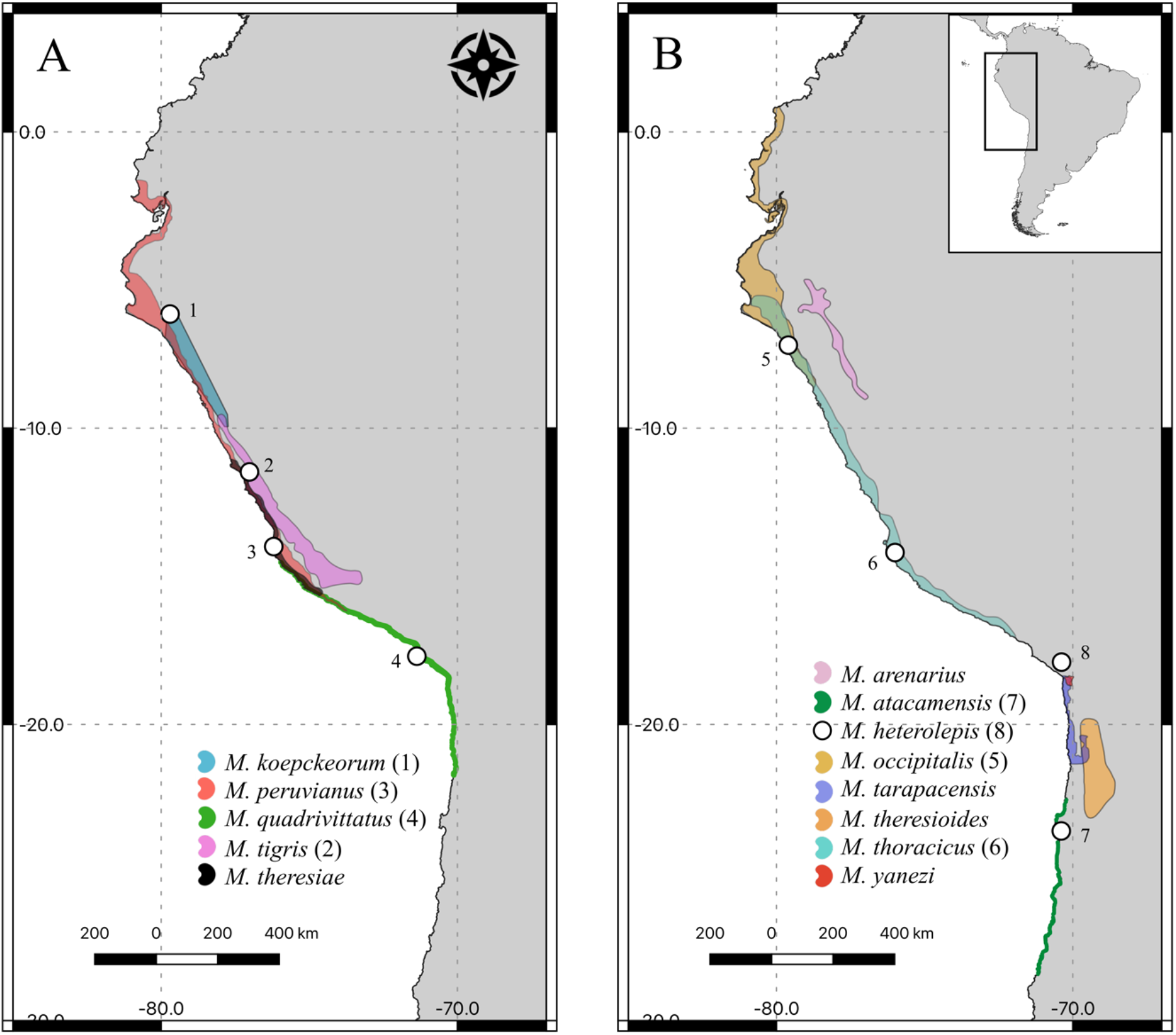
Ranges of continental *Microlophus* species. Distributions for each species are represented by different colors as shown in the legend; taxa are split across two plots for clarity. For the seven species with the broadest distributions, points (1-7, also shown in the legend) represent the localities from which maximum body size data and climatic variables were obtained. For the rest of species, the average climatic variables across their distributions and maximum body sizes reported in the literature were considered (see Methods). The average of the only two locality coordinates known for *M. heterolepis*, is represented by point number 8.

As mentioned above, previous research suggests that latitudinal and climatic gradients of body size are highly variable, especially among ectotherms, and that the mechanisms driving these associations differ among taxa and ecological contexts. Furthermore, the relationships between body size and climatic conditions might help explain other macroevolutionary patterns that involve size. The vast latitudinal distribution of the genus *Microlophus*, paired with the variability of body sizes and levels of sexual size dimorphism shown by its species, provide an interesting opportunity to test for Bergmann’s rule as well as examine the mechanisms that underlie it, and the effects it might have on patterns of size dimorphism. Therefore, in this study we (1) test for an association between body size and latitude in *Microlophus* lizards, (2) test whether this association is different between sexes, potentially producing a latitudinal gradient of sexual size dimorphism, and (3) explore possible mechanisms for these relationships by testing for associations between body size and climatic variables.

## METHODS

### Body size data

We used body size data recently compiled by Toyama and Boccia (2021). These data are based on maximum SVLs reported in the literature for males and females for each *Microlophus* species. Some of these species occur across a wide range of latitudes, with different populations potentially showing different body sizes along the latitudinal gradient (Figure 1, see also Dixon and Wright, 1975; Sepúlveda et al., 2014). Thus, the maximum reported body sizes for each species might not correspond to the average latitude of their distributions. To avoid this mismatch, we considered the largest reported sizes for males and females for the seven species with the broadest latitudinal distributions (*M. atacamensis, M. koepckeorum, M. occipitalis, M. peruvianus, M. quadrivittatus, M. thoracicus, M. tigris*) for which geographic coordinates corresponding to data collection localities were available and used the climatic data corresponding to those coordinates. For the rest of species, we extracted the climatic data as described in the next section and used the maximum body sizes reported in the literature. All body size data were log-transformed.

### Climatic data

We extracted distributional data from the International Union for the Conservation of Nature (IUCN) repository for 21 species for which these data were available. For the remaining 2 species we obtained point coordinates from GBIF (*M. barringtonensis*) or used literature locality data (*M. heterolepis*). Since GBIF draws from multiple data sources, we cross-checked all point coordinates against known localities for *M. barringtonensis* and removed any erroneous observations. Occurrence points for *M. heterolepis* were taken from Vidal et al. (2017). We downloaded climatic data corresponding to the geographic distributions of all species from WorldClim (0.5 arc-minute resolution) (Fick and Hijmans, 2017). Data rasters were assembled using *WorldClimTiles* and *raster* R packages (Kapitza, 2019; Hijmans et al, 2015). We considered all 19 climatic variables available and two geographic variables: altitude and latitude. Climatic variables included annual, quarterly, and monthly mean temperatures and precipitation levels (selected quarters or months only, see Tables 1 and 2), diurnal temperature range, temperature and precipitation seasonality (measures of temperature and precipitation changes over the course of a year), and isothermality (which quantifies how large the day-to-night temperatures oscillate relative to the summer-to-winter (annual) oscillations). Precipitation variables were log-transformed given the skewed distribution of this type of data. Range-wide means for each variable were calculated for each polygon or point set corresponding to a species. Additionally, as mentioned in the previous section, climatic data was extracted from morphological data set collection coordinates for the seven species with the broadest latitudinal distributions. The R script and resulting data files are available at https://github.com/chrisboccia/microlophus_bergmann/.

**Table 1.**
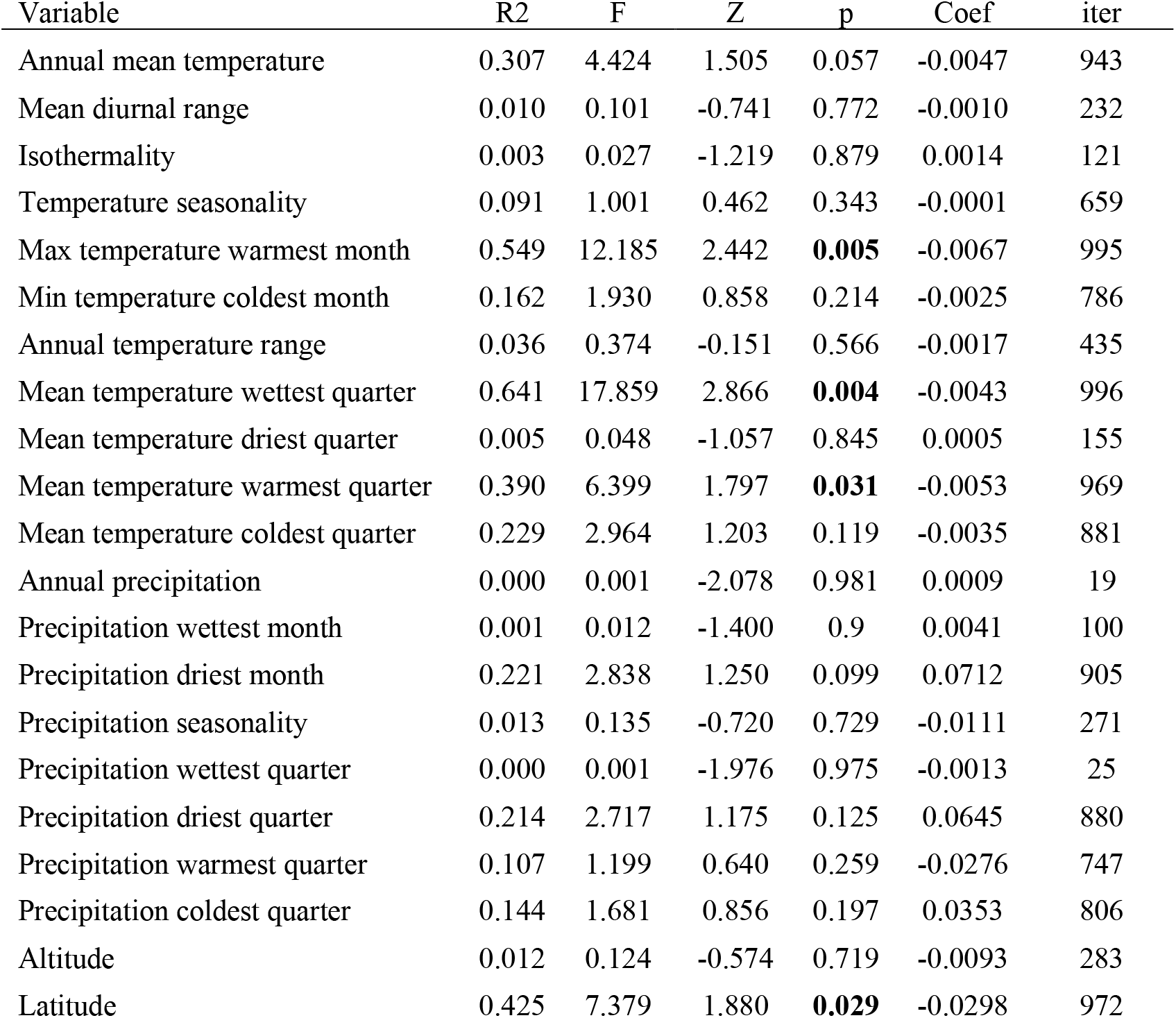
Mean values of ANOVA statistics from pairwise PGLS regressions between climatic variables and male body size in mainland *Microlophus*. Significant p-values are shown in bold. The estimated value for the regression coefficient (Coef) and the number of cases that do not exceed the magnitude of the observed coefficient over 1000 phylogeny iterations (iter) are also provided.

**Table 2.** Mean values of ANOVA statistics from pairwise PGLS regressions between climatic variables and female body size in mainland *Microlophus*. Significant p-values are shown in bold. The estimated value for the regression coefficient (Coef) and the number of cases that do not exceed the magnitude of the observed coefficient over 1000 phylogeny iterations (iter) are also provided.

| Variable | R2 | F | Z | p | Coef | iter |
| --- | --- | --- | --- | --- | --- | --- |
| Annual mean temperature | 0.338 | 5.106 | 1.633 | **0.043** | -0.0044 | 957 |
| Mean diurnal range | 0.007 | 0.071 | -0.832 | 0.795 | -0.0007 | 207 |
| Isothermality | 0.002 | 0.019 | -1.223 | 0.877 | 0.0011 | 123 |
| Temperature seasonality | 0.040 | 0.414 | -0.100 | 0.548 | -0.0001 | 459 |
| Max temperature warmest month | 0.604 | 15.283 | 2.660 | **0.004** | -0.0064 | 996 |
| Min temperature coldest month | 0.167 | 2.011 | 0.891 | 0.199 | -0.0023 | 802 |
| Annual temperature range | 0.027 | 0.276 | -0.298 | 0.629 | -0.0012 | 373 |
| Mean temperature wettest quarter | 0.495 | 9.805 | 2.275 | **0.008** | -0.0034 | 992 |
| Mean temperature driest quarter | 0.002 | 0.021 | -1.289 | 0.888 | -0.0003 | 116 |
| Mean temperature warmest quarter | 0.405 | 6.804 | 1.865 | **0.031** | -0.0048 | 969 |
| Mean temperature coldest quarter | 0.282 | 3.934 | 1.421 | 0.069 | -0.0035 | 931 |
| Annual precipitation | 0.008 | 0.083 | -0.800 | 0.777 | -0.0079 | 225 |
| Precipitation wettest month | 0.000 | 0.002 | -1.889 | 0.973 | 0.0013 | 27 |
| Precipitation driest month | 0.125 | 1.430 | 0.699 | 0.279 | 0.0487 | 727 |
| Precipitation seasonality | 0.038 | 0.398 | -0.108 | 0.533 | -0.0157 | 471 |
| Precipitation wettest quarter | 0.006 | 0.063 | -0.908 | 0.794 | -0.0076 | 208 |
| Precipitation driest quarter | 0.055 | 0.588 | 0.100 | 0.472 | 0.0259 | 533 |
| Precipitation warmest quarter | 0.094 | 1.041 | 0.560 | 0.305 | -0.0230 | 700 |
| Precipitation coldest quarter | 0.006 | 0.059 | -0.927 | 0.819 | 0.0062 | 182 |
| Altitude | 0.025 | 0.251 | -0.300 | 0.607 | -0.0114 | 396 |
| Latitude | 0.285 | 3.991 | 1.429 | 0.073 | -0.0217 | 930 |

### Analyses

We performed analyses on three different datasets that considered (1) all species, (2) only mainland species, and (3) only mainland species without *M. arenarius*. We decided to remove island species from the latter pair of datasets because of their limited latitudinal distribution and because unique insular ecological dynamics might affect the evolution of body size on islands (Andrews, 1976; Palkovacs, 2003). We also removed *M. arenarius* from one set of analyses because of its unusual distribution (Figure 1B). To account for phylogenetic relatedness in our analyses, a phylogenetic hypothesis for *Microlophus* was trimmed from the squamate phylogeny of Pyron et al. (2013). The trimmed phylogeny included 19 of the 23 *Microlophus* species (*M. barringtonensis, M. indefatigabilis, M. jacobii* and *M. tarapacensis* were not present in the original phylogeny).

To test for a pattern consistent with Bergmann’s rule, we performed phylogenetic generalized least squares (PGLS) regressions (function “gls” from the R package *nlme*, Pinheiro et al. 2018) between log-transformed body size and latitude (southern latitudes being negative numbers) for each of the three datasets, and for males and females separately. A negative relationship between size and latitude obtained from these tests would support Bergmann’s rule. Posteriorly, and only for mainland species (dataset 2), we tested whether the relationship between size and latitude was different between sexes comparing the absolute difference between the slopes of PGLS regressions (size∼latitude) performed on each sex separately against a distribution of absolute differences obtained from a permutation analysis (1000 iterations).

Finally, in order to uncover the possible mechanisms behind latitude-body size associations, we performed pairwise PGLS regressions between body size and each of the range descriptors (i.e., geographic and climatic range variables). The significance of all PGLS models was evaluated using residual randomization procedures implemented in the R package *RRPP* (Collyer and Adams, 2018). All the regressions considered a covariance matrix based on the estimation of Pagel’s lambda. These regressions were performed for the three species datasets and for males and females separately. We focus on the results obtained from dataset 2 in the main text, but results for the other datasets are detailed in the supplemental materials. All analyses were performed in R (R Core Team 2021).

## RESULTS

### Bergmann’s rule in Microlophus and sex-specific latitudinal gradients

In accordance with Bergmann’s rule, we found a negative relationship between body size and latitude across both sexes in *Microlophus* lizards. Although this relationship was not significant when considering all species (Tables S1, S2), it was supported when considering only mainland species (Figure 2A,B; Tables 1, 2). Removing *M. arenarius* from the analysis resulted in an even stronger negative relationship (Figure 2A,B, Tables S3, S4). The slope of the relationship between body size and latitude was steeper in males than in females among *Microlophus* species (Figure 3). However, based on a permutation analysis, the slopes of these relationships were not significantly different (*p* = 0.539; Figure 3).

**Figure 2.**
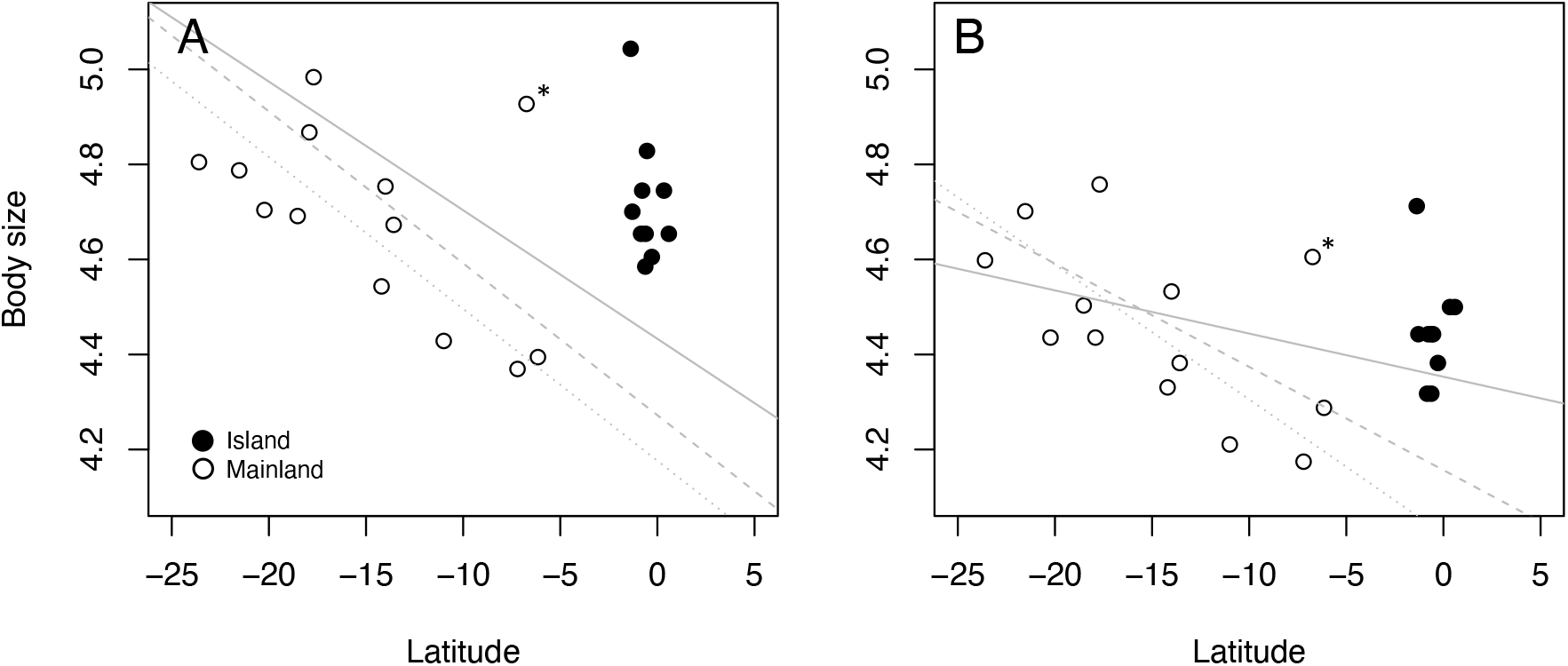
Relationships between body size and latitude for (A) male and (B) female *Microlophus*. Closed and open circles represent island and mainland species, respectively. The asterisks indicate the position of *M. arenarius*. Straight lines represent the fitted PGLS regression models considering all species (continuous line), mainland species (dashed line) and mainland species without *M. arenarius* (dotted line).

**Figure 3.**
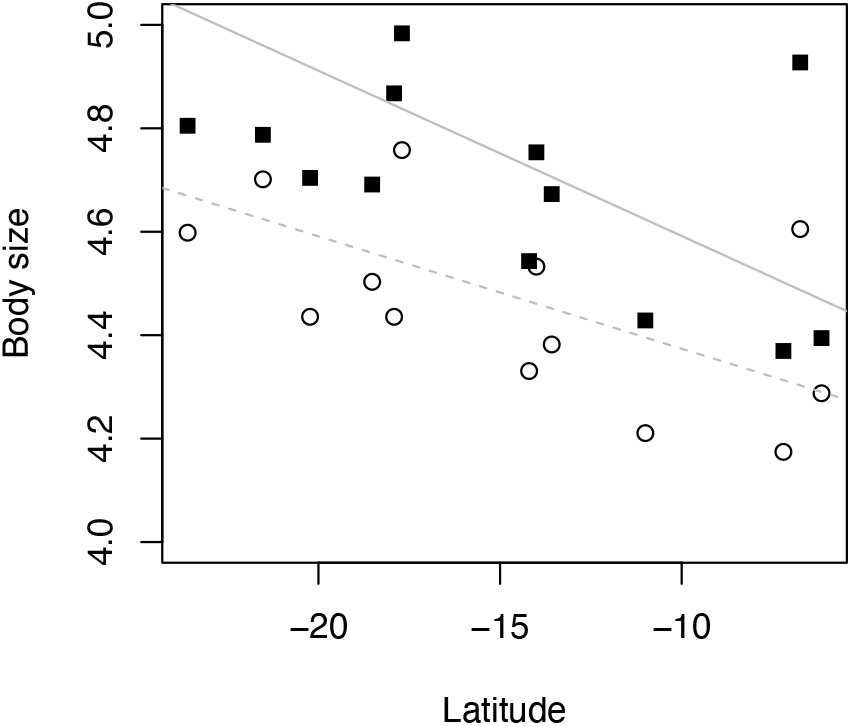
Relationship between body size and latitude for males (black squares, continuous line) and females (white circles, dashed line) of mainland species. Lines represent the fitted PGLS regression models. No significant differences in slopes were found between sexes (permutation analysis, *p* = 0.539).

### Association between body size and climatic variables

Pairwise PGLS analyses between body size and climatic variables considering only mainland species identified the following associations. In males, body size was significantly and negatively related to three temperature variables: maximum temperature of the warmest month and mean temperatures of the wettest and warmest quarters (Table 1). Similar relationships were found for females, with the addition of a negative relationship between size and annual mean temperature (Table 2).

Temperature-related variables were still strongly related to size when removing *M. arenarius*, but in this case the negative effects of precipitation and isothermality were also detected (Tables S3, S4). Considering all species, we identified a negative relationship between body size and maximum temperature of the warmest month for both males and females (Tables S1, S2). In summary, larger body sizes in continental *Microlophus* occur at southern latitudes, and are associated with colder, drier, and more thermally variable environments.

## DISCUSSION

### Bergmann’s rule: pattern

The relationship we observed between body size and latitude in *Microlophus* was consistent with Bergmann’s rule, especially among mainland species. Larger species were found at higher latitudes than smaller ones (Figure 2). This finding is consistent with previous research in other squamates (e.g., Ashton and Feldman, 2003; Cruz et al., 2005) though it contrasts with work on other groups of lizards (Pincheira-Donoso et al., 2008).

Latitudinal patterns of body size were similar for males and females (Figure 3). Blanckenhorn et al (2006) showed that, at the intraspecific level, a latitudinal version of Rensch’s rule (a positive relationship between sexual size dimorphism and species size) arose in a majority of taxa as a consequence of males and females showing different associations between body size and latitude. Similarly, Tarr et al (2019) showed that Central and North American lizard species exhibited an increase in male-biased sexual size dimorphism towards lower latitudes which was driven by a positive association between male body size and temperature. This strong association was explained by a “reproductive selection hypothesis” which states that females lay more clutches throughout the year in aseasonal habitats, resulting in stronger sexual selection on males as the benefits of controlling territories are higher when reproduction is continuous (Machado et al., 2016). Our results showed that the body size-latitude relationships were not different between males and females, indicating that patterns of sexual size dimorphism in *Microlophus* are not being driven by different latitudinal gradients of male and female body size and support previous results indicating no evidence of Rensch’s rule in *Microlophus* lizards (Toyama and Boccia, 2021). Additionally, our results suggest that the latitudinal gradients of body size in *Microlophus* probably do not depend on latitudinal gradients of sexual selection acting on males as expected based on the “reproductive selection hypothesis”. In fact, the relationship between size and latitude found for *Microlophus* species was opposite to the one found by Tarr et al for other lizard species.

*M. arenarius* is much larger in size than the other continental *Microlophus* after accounting for the effect of latitude (Figure 2). *Microlophus arenarius* is the only species of the genus not found in arid coastal regions, suggesting that its relatively large size could be a consequence of its unique ecology. For example, the inter-Andean valleys where *M. arenarius* is found receive a higher amount of precipitation throughout the year when compared to the coastal arid regions, where rainfall is almost non-existent (Duellman and Pramuk, 1999; Rundel et al., 2007). Higher precipitation is related to increased primary productivity, which in turn increases the availability of resources and could allow the maintenance of larger body sizes (Yom-Tov and Geffen, 2006). Moreover, the diet of *M. arenarius* includes large quantities of plant material (Beuttner and Koch, 2019), which is readily available in the highly vegetated areas where the species is found as a consequence of regular precipitation. Large body sizes and a warm environment have been traditionally considered as essential requirements for herbivory in lizards due to the physiological difficulties of plant digestion (Pough, 1973; Cooper and Vitt, 2002), suggesting that the large size of *M, arenarius* could be an adaptation for plant consumption. However, there is strong evidence against an obligate relationship between body size and herbivory (Espinoza et al., 2004; Vitt, 2004), including the fact that other *Microlophus* species that include large proportions of plant material in their diets (e.g., *M. koepckeorum* and *M. thoracicus*, Pérez et al., 2015; Toyama et al., 2018; Pérez, unpublished data) are not among the largest in size within the genus and do not inhabit particularly warm environments. A closer look at the ecology of *M. arenarius* and a comparison with the rest of continental species could provide more detailed insights regarding its large size.

Insular species are also larger than expected for the latitudinal range at which they are found (Figure 2). Ecological differences between island and mainland environments can result in the evolution of different life history traits between insular and continental species (i.e., the “island syndrome”, Adler and Levins, 1994; Novosolov et al., 2013), potentially affecting the evolution of body size. For example, the lack of predators on islands can delay maturity and favour long term intraspecific competition, resulting in larger adult sizes. On the other hand, depauperate islands might be poor in food resources compared to mainland habitats, resulting in slower growth rates and smaller body sizes (Palkovacs, 2003). These dynamics can result in what is termed the “island rule”, the tendency of small and large taxa to exhibit trends towards gigantism and dwarfism on islands, respectively (Meiri, 2007). A previous comparison between island and mainland *Microlophus* species showed no differences between the average body sizes of these two groups (Toyama and Boccia, 2021), contradicting the expectations of the island rule. However, this could just mean that this comparison is overlooking the effect of latitude. For example, in a large-scale study on lizards, Stark et al (2018) found that life spans and, to some extent, body size, were positively related to insularity, seasonality, and low temperatures, supporting the possibility of an interaction between insularity and latitude in producing body size clines. *Microlophus* species from the Galápagos are larger than mainland species found at similar latitudes, but it remains unclear whether insular species might get gradually larger at higher latitudes. With little variation in insular latitude however, comparisons of latitudinal gradients of body size between insular and mainland species are not possible in *Microlophus*. Future studies considering island-mainland systems with large insular latitudinal ranges could help test this hypothesis.

### Bergmann’s rule: mechanism

Body size was significantly and negatively related to temperature variables, as expected based on the heat conservation mechanism proposed by Bergmann (Meiri, 2011). Traditionally, ectotherms are not expected to adhere to this mechanism, as they do not produce heat internally and the costs of gaining heat would exceed the benefits of conserving it at cold climates and large body sizes (Cushman et al., 1993). However, despite this expectation some studies suggest that ectotherms can still benefit from this mechanism. For example, Zamora-Camacho et al (2014) found that the body size of *Psammodromus algirus* lizards increased with altitude, and while their cooling rates decreased with size as a result of thermal inertia, the heating rates did not vary significantly across the altitudinal gradient, meaning that larger lizards were more efficient in conserving heat in colder environments without significant costs related to heat gain. This suggests that increasing body size in cooler climates might actually have a positive effect in maintaining a preferred body temperature in some ectotherms, supporting Bergmann’s original mechanism. Thermoregulatory dynamics, however, might be specific to particular taxa. In contrast with the results of Zamora-Camacho et al. (2014) for *P. algirus*, it is known that in *M. atacamensis* larger sizes are related to lower cooling rates, but also to lower heating rates (Sepúlveda et al., 2008) representing a difficulty for the heat conservation hypothesis. More tests of this kind in other *Microlophus* species could help determine whether a negative association between size and temperature could be explained by the traditional heat conservation mechanism or whether other mechanisms might play a more important role.

When we removed *M. arenarius* from the analyses some precipitation variables and isothermality appeared to be negatively related to body size (Tables S3, S4). This is unexpected given that precipitation is usually positively related with body size because of its association with primary production (Yom-Tov and Geffen, 2006). However, precipitation variables might be inadequate to interpret the observed latitudinal patterns given that most *Microlophus* species inhabit regions with high aridity and negligible levels of rainfall (Rundel et al., 2007). Isothermality, which quantifies the ratio of day-to-night temperature oscillations to summer-to-winter (annual) oscillations, might be an indicator of seasonality across latitudes (MacArthur, 1972). Besides its relationship to the reproductive selection hypothesis discussed above, seasonality has been identified as a possible driver of latitudinal gradients of body size through its effect on resource availability. As habitats become more seasonal at higher latitudes, food shortages also become more frequent (Boyce, 1978). In this context, larger sizes would be beneficial at high latitudes because they allow individuals to survive for long periods of time without food, in what is known as the “starvation hypothesis” (Morrison, 1960; Lindstedt and Boyce, 1985; Arnett and Gotelli, 2003). Although *M. arenarius* is a large species, it is distributed in habitats with low seasonality (Duellman and Pramuk, 1999), suggesting that its large size is not an adaptation for long periods of starvation. For the rest of the continental species, the availability of trophic resources throughout the year and its relationship with latitude need to be quantified to test this hypothesis. *Microlophus* species have diverse diets, and are known to consume arthropods, plant material, algae, and crustaceans; they have even been observed to foray into cannibalism (Pérez and Balta, 2005, 2007; Quispitúpac and Pérez, 2008; Sepúlveda et al., 2014; Toyama et al., 2018). However, it is so far unknown whether the species’ diets vary latitudinally and whether the availability of some dietary items are more affected by seasonality than others.

Our results provide evidence for a latitudinal gradient of body size in *Microlophus* lizards consistent with Bergmann’s rule; this gradient is especially strong when we considered only mainland species. We also provide evidence for an effect of temperature, precipitation, and seasonality in driving this pattern. We discussed how the lack of sexual differences in latitudinal gradients of body size is consistent with what is known about sexual size dimorphism in *Microlophus* and how the relationships between body size and climatic variables could support some potential mechanisms driving the observed body size gradient. The vast latitudinal gradient represented by the geographical distribution of *Microlophus* lizards provides an exceptional opportunity to evaluate the effects of climatic and ecological variables on the evolution of body size. However, our knowledge of this system is still limited, and future ecological and natural history studies will be vital if we want to increase our understanding of the mechanisms driving the patterns we observed.

## ACKNOWLEDGMENTS

We thank Miriam Ahmad-Gawel for useful comments, corrections, and suggestions on previous versions of this manuscript.

