## Supplemental Materials for "Bergmann’s rule in *Microlophus* lizards: testing for latitudinal and climatic gradients of body size"

Table S1. Mean values of ANOVA statistics from pairwise PGLS regressions between climatic variables and male body size in all *Microlophus* species (including both island and mainland species). Significant p-values are shown in bold. The estimated value for the regression coefficient (Coef) and the number of cases that do not exceed the magnitude of the observed coefficient over 1000 phylogeny iterations (iter) are also provided.

| Variable | R2 | F | Z | p | Coef | iter |
|---|---|---|---|---|---|---|
| Annual mean temperature | 0.026 | 0.460 | -0.010 | 0.51 | -0.0014 | 490 |
| Mean diurnal range | 0.071 | 1.291 | 0.662 | 0.272 | -0.0022 | 729 |
| Isothermality | 0.024 | 0.420 | -0.049 | 0.537 | -0.0037 | 463 |
| Temperature seasonality | 0.067 | 1.220 | 0.618 | 0.297 | -0.0001 | 703 |
| Max temperature warmest month | 0.251 | 5.686 | 1.789 | **0.033** | -0.0052 | 967 |
| Min temperature coldest month | 0.000 | 0.000 | -2.532 | 0.997 | 0.0000 | 3 |
| Annual temperature range | 0.090 | 1.681 | 0.847 | 0.209 | -0.0022 | 792 |
| Mean temperature wettest quarter | 0.213 | 4.599 | 1.538 | 0.065 | -0.0030 | 937 |
| Mean temperature driest quarter | 0.001 | 0.015 | -1.379 | 0.909 | 0.0002 | 91 |
| Mean temperature warmest quarter | 0.081 | 1.492 | 0.753 | 0.242 | -0.0023 | 758 |
| Mean temperature coldest quarter | 0.011 | 0.182 | -0.498 | 0.703 | -0.0008 | 298 |
| Annual precipitation | 0.019 | 0.337 | -0.133 | 0.563 | 0.0138 | 440 |
| Precipitation wettest month | 0.023 | 0.405 | -0.039 | 0.528 | 0.0186 | 472 |
| Precipitation driest month | 0.174 | 3.571 | 1.459 | 0.063 | 0.0450 | 938 |
| Precipitation seasonality | 0.009 | 0.153 | -0.384 | 0.62 | -0.0115 | 380 |
| Precipitation wettest quarter | 0.016 | 0.273 | -0.244 | 0.613 | 0.0140 | 387 |
| Precipitation driest quarter | 0.177 | 3.655 | 1.504 | 0.062 | 0.0438 | 939 |
| Precipitation warmest quarter | 0.024 | 0.412 | -0.025 | 0.511 | -0.0139 | 490 |
| Precipitation coldest quarter | 0.142 | 2.817 | 1.295 | 0.093 | 0.0301 | 907 |
| Altitude | 0.011 | 0.189 | -0.465 | 0.688 | -0.0118 | 312 |
| Latitude | 0.163 | 3.318 | 1.423 | 0.071 | -0.0189 | 930 |

Table S2. Mean values of ANOVA statistics from pairwise PGLS regressions between climatic variables and female body size in all *Microlophus* species (including both island and mainland species). Significant p-values are shown in bold. The estimated value for the regression coefficient (Coef) and the number of cases that do not exceed the magnitude of the observed coefficient over 1000 phylogeny iterations (iter) are also provided.

| Variable | R2 | F | Z | p | Coef | iter |
|---|---|---|---|---|---|---|
| Annual mean temperature | 0.037 | 0.652 | 0.202 | 0.443 | -0.0015 | 559 |
| Mean diurnal range | 0.029 | 0.506 | 0.024 | 0.501 | -0.0012 | 500 |
| Isothermality | 0.001 | 0.021 | -1.294 | 0.889 | -0.0008 | 111 |
| Temperature seasonality | 0.042 | 0.752 | 0.281 | 0.42 | -0.0001 | 581 |
| Max temperature warmest month | 0.308 | 7.584 | 1.972 | **0.019** | -0.0049 | 981 |
| Min temperature coldest month | 0.005 | 0.080 | -0.822 | 0.786 | -0.0003 | 214 |
| Annual temperature range | 0.051 | 0.915 | 0.398 | 0.372 | -0.0014 | 630 |
| Mean temperature wettest quarter | 0.153 | 3.079 | 1.250 | 0.115 | -0.0019 | 885 |
| Mean temperature driest quarter | 0.002 | 0.029 | -1.154 | 0.862 | 0.0003 | 138 |
| Mean temperature warmest quarter | 0.106 | 2.008 | 0.950 | 0.188 | -0.0024 | 813 |
| Mean temperature coldest quarter | 0.024 | 0.423 | -0.048 | 0.537 | -0.0011 | 466 |
| Annual precipitation | 0.000 | 0.001 | -1.834 | 0.963 | 0.0007 | 37 |
| Precipitation wettest month | 0.006 | 0.110 | -0.634 | 0.721 | 0.0085 | 280 |
| Precipitation driest month | 0.096 | 1.797 | 0.941 | 0.172 | 0.0296 | 828 |
| Precipitation seasonality | 0.034 | 0.590 | 0.219 | 0.424 | -0.0162 | 577 |
| Precipitation wettest quarter | 0.000 | 0.004 | -1.596 | 0.93 | 0.0015 | 70 |
| Precipitation driest quarter | 0.062 | 1.133 | 0.633 | 0.277 | 0.0210 | 723 |
| Precipitation warmest quarter | 0.030 | 0.526 | 0.131 | 0.461 | -0.0133 | 540 |
| Precipitation coldest quarter | 0.020 | 0.355 | -0.076 | 0.542 | 0.0101 | 459 |
| Altitude | 0.041 | 0.719 | 0.263 | 0.408 | -0.0163 | 592 |
| Latitude | 0.056 | 1.006 | 0.525 | 0.308 | -0.0091 | 694 |

Table S3. Mean values of ANOVA statistics from pairwise PGLS regressions between climatic variables and male body size in mainland *Microlophus* species (not including *M. arenarius*). Significant p-values are shown in bold. The estimated value for the regression coefficient (Coef) and the number of cases that do not exceed the magnitude of the observed coefficient over 1000 phylogeny iterations (iter) are also provided.

| Variable | R2 | F | Z | p | Coef | iter |
|---|---|---|---|---|---|---|
| Annual mean temperature | 0.310 | 4.036 | 1.483 | 0.065 | -0.0044 | 935 |
| Mean diurnal range | 0.080 | 0.782 | 0.263 | 0.406 | -0.0026 | 598 |
| Isothermality | 0.480 | 8.308 | 1.876 | **0.028** | -0.0262 | 972 |
| Temperature seasonality | 0.077 | 0.751 | 0.324 | 0.376 | 0.0002 | 631 |
| Max temperature warmest month | 0.500 | 9.000 | 2.078 | **0.014** | -0.0063 | 986 |
| Min temperature coldest month | 0.035 | 0.330 | -0.150 | 0.583 | -0.0010 | 425 |
| Annual temperature range | 0.015 | 0.135 | -0.562 | 0.699 | -0.0009 | 309 |
| Mean temperature wettest quarter | 0.674 | 18.626 | 2.765 | **0.004** | -0.0041 | 996 |
| Mean temperature driest quarter | 0.054 | 0.516 | 0.076 | 0.485 | 0.0013 | 522 |
| Mean temperature warmest quarter | 0.312 | 4.082 | 1.488 | 0.066 | -0.0048 | 934 |
| Mean temperature coldest quarter | 0.319 | 4.217 | 1.520 | 0.059 | -0.0038 | 941 |
| Annual precipitation | 0.097 | 0.964 | 0.380 | 0.378 | -0.0321 | 627 |
| Precipitation wettest month | 0.062 | 0.597 | 0.120 | 0.482 | -0.0296 | 525 |
| Precipitation driest month | 0.000 | 0.000 | -2.246 | 0.989 | 0.0088 | 11 |
| Precipitation seasonality | 0.013 | 0.116 | -0.713 | 0.75 | -0.0093 | 253 |
| Precipitation wettest quarter | 0.095 | 0.944 | 0.382 | 0.386 | -0.0339 | 622 |
| Precipitation driest quarter | 0.033 | 0.311 | -0.235 | 0.592 | 0.0359 | 412 |
| Precipitation warmest quarter | 0.516 | 9.583 | 2.003 | **0.019** | -0.0640 | 982 |
| Precipitation coldest quarter | 0.011 | 0.098 | -0.662 | 0.735 | 0.0101 | 267 |
| Altitude | 0.083 | 0.814 | 0.266 | 0.404 | -0.0215 | 606 |
| Latitude | 0.573 | 12.059 | 2.342 | **0.009** | -0.0325 | 992 |

Table S4. Mean values of ANOVA statistics from pairwise PGLS regressions between climatic variables and female body size in mainland *Microlophus* species (not including *M. arenarius*). Significant p-values are shown in bold. The estimated value for the regression coefficient (Coef) and the number of cases that do not exceed the magnitude of the observed coefficient over 1000 phylogeny iterations (iter) are also provided.

| Variable | R2 | F | Z | p | Coef | iter |
|---|---|---|---|---|---|---|
| Annual mean temperature | 0.238 | 2.811 | 1.193 | 0.11 | -0.0037 | 892 |
| Mean diurnal range | 0.042 | 0.394 | -0.099 | 0.544 | -0.0017 | 465 |
| Isothermality | 0.311 | 4.068 | 1.301 | 0.09 | -0.0198 | 910 |
| Temperature seasonality | 0.069 | 0.663 | 0.214 | 0.423 | 0.0002 | 581 |
| Max temperature warmest month | 0.580 | 12.422 | 2.369 | **0.008** | -0.0064 | 992 |
| Min temperature coldest month | 0.054 | 0.519 | 0.104 | 0.475 | -0.0012 | 537 |
| Annual temperature range | 0.010 | 0.093 | -0.783 | 0.782 | -0.0007 | 220 |
| Mean temperature wettest quarter | 0.492 | 8.729 | 2.039 | **0.015** | -0.0033 | 985 |
| Mean temperature driest quarter | 0.003 | 0.025 | -1.192 | 0.863 | 0.0003 | 138 |
| Mean temperature warmest quarter | 0.368 | 5.248 | 1.665 | **0.041** | -0.0049 | 959 |
| Mean temperature coldest quarter | 0.242 | 2.871 | 1.207 | 0.108 | -0.0032 | 893 |
| Annual precipitation | 0.098 | 0.978 | 0.391 | 0.368 | -0.0291 | 638 |
| Precipitation wettest month | 0.030 | 0.278 | -0.253 | 0.593 | -0.0188 | 412 |
| Precipitation driest month | 0.014 | 0.132 | -0.573 | 0.772 | -0.7698 | 250 |
| Precipitation seasonality | 0.042 | 0.394 | -0.087 | 0.509 | -0.0151 | 494 |
| Precipitation wettest quarter | 0.077 | 0.752 | 0.248 | 0.424 | -0.0277 | 586 |
| Precipitation driest quarter | 0.014 | 0.125 | -0.601 | 0.722 | -0.0206 | 282 |
| Precipitation warmest quarter | 0.306 | 3.971 | 1.288 | 0.092 | -0.0440 | 911 |
| Precipitation coldest quarter | 0.429 | 6.768 | 1.810 | **0.026** | -0.0466 | 976 |
| Altitude | 0.078 | 0.757 | 0.241 | 0.423 | -0.0192 | 590 |
| Latitude | 0.653 | 16.973 | 2.674 | **0.003** | -0.0281 | 997 |
